# Promoter- and Enhancer-Dependent Cohesin Loading Initiates Chromosome Looping to Fold *Tcrb* Loci for Long-Range Recombination

**DOI:** 10.1101/2024.10.13.618029

**Authors:** Brittney M. Allyn, Katharina E. Hayer, Clement Oyeniran, Vincent Nganga, Kyutae Lee, Ahmet Sacan, Eugene M. Oltz, Craig H. Bassing

## Abstract

Cohesin-mediated chromosome looping regulates diverse processes, including antigen receptor (AgR) gene assembly by V(D)J recombination. To understand mechanisms that coordinate genome topologies, we focused on a genetically tractable AgR locus, *Tcrb*. Cohesin loading and initiating loop extrusion (LE) from a nearby CTCF-binding element (CBE) required the promoter of the most 5’Vβ segment, creating long-range contacts with target downstream DJβ segments within the recombination center (RC). CBEs flanking the RC have multiple functions: terminators of LE originating in the Vβ cluster, initiators of LE in the RC, and insulation of enhancer activity. Deletion of the *Tcrb* super-enhancer abolished loop extrusion from the neighboring RC but spared long-range contacts, indicating that unidirectional loop extrusion from upstream Vβ segments was sufficient. Thus, Vβ promoter- or enhancer-dependent cohesin loading initiates LE in opposite directions across the locus to assemble a broad *Tcrb* repertoire, a finding that has broad implications for genomic architecture and function.

## Introduction

The dynamic organization of genome topology has diverse fundamental roles in DNA transcription, replication, segregation, repair, and recombination. Two general mechanisms shape genome topologies: homotypic chromatin interactions between regions of similar transcriptional activity and point-to-point chromosome looping. The former mechanism encompasses a collection of chromatin-based processes, including phase separation of DNA binding proteins, interactions between similar histone modifications, and formation of transcription factories^1–6^. The latter involves creation of loops by the cohesin protein complex^7,8^, which loads onto DNA and subsequently extrudes or translocates along DNA, processes each assisted by the NIPBL protein^9,10^. When cohesin encounters an impediment, it reels DNA from the unblocked side and drives loop extrusion until encountering another obstacle, resulting in a chromosome loop^11–13^. Most chromosomal loops are anchored by a pair of convergently oriented CTCF binding elements (CBEs), which function as significant impediments to continued cohesin-mediated extrusion^8,14,15^. However, chromosome loops can occur between CBEs of the same orientation, or a CBE and another impediment, such as active transcription or other chromatin bound proteins^16–19^. Correlations between transcriptional activation and cohesin localization imply mechanistic relationships between homotypic chromatin interactions and cohesin/CTCF-directed chromosome looping. In addition, transcription of non-coding RNAs can regulate genome topology through homotypic interactions or as impediments for chromosome looping. Beyond serving as anchors for loops, CBEs also can function as insulators that block promoter-enhancer interactions and/or the spread of active and inactive chromatin, with these functions also regulating gene transcription and genome topology^20–23^. The growing number of disease-associated mutations that alter genome topology^24^ has heightened interest in understanding molecular mechanisms that fold genomes to control gene expression levels and to restrict DNA translocations.

An important biological process in which the dynamic regulation of genome topology governs beneficial versus pathological outcomes is diversification of *Igh*, *Igk*, *Tcra/d*, and *Tcrb* lymphocyte antigen receptor (AgR) genes via recombination of variable (V), diversity (D), and joining (J) gene segments across linear genomic distances as large as 3 Mb^25–30^. The RAG endonuclease initiates AgR gene assembly by cleaving at recombination signal sequence (RSS) elements that flank a pair of gene segments, with subsequent repair of these DNA breaks by the non-homologous end joining (NHEJ) pathway to form a chromosomal coding join between the gene segments^31,32^. The rearranged V(D)J segments forms the variable region exon, which together with downstream constant (C) region exons, comprises an assembled AgR gene. Each AgR locus is activated for transcription in the lymphocyte lineage and at the developmental stage in which they recombine. This creates a chromatin landscape that promotes RAG binding over the (D)J portion of an AgR locus, forming the aptly named recombination center (RC)^33^. Independent of RAG binding, AgR loci undergo changes in topology that compact (or fold) them, juxtaposing their V segments and RCs to facilitate RAG-mediated synapsis and cleavage of RSSs across vast linear genomic distances^30,34–39^. It has long been appreciated that AgR promoters and enhancers render gene segments accessible for RAG binding and synapsis; yet, little is known about the functions of these *cis*-acting elements or resultant non-coding RNAs (ncRNAs) in the process of locus folding. In this context, while homotypic interactions between transcriptionally active V and (D)J chromatin have been proposed to fold loci and facilitate long-range recombination^40^, the sole experimental evidence for this derives from *Tcrb* ^39^. In contrast, the roles of chromosome looping in long-range V(D)J recombination is now supported by numerous studies revealing that cohesin, CTCF, and/or CBEs promote locus compaction to enhance interactions between V segments and RCs at *Igh*, *Igk*, *Tcra/d*, and *Tcrb*^19–22,39,41–43^. However, conformational analyses indicate that each AgR locus likely adopts unique patterns of long-range looping, which could provide insights into the range of mechanisms employed throughout the genome for establishing and revising architectural domains.

At *Igh* and *Tcrd*, cohesin-mediated loop extrusion predominantly, if not exclusively, proceeds from downstream of the RC and tracks upstream through V chromatin. This directionality enables RAG bound at a D RSS to linearly and uni-directionally scan upstream DNA in search of an accessible V RSS of convergent genomic orientation for synapsis and deletional recombination^19,25,44–48^. In contrast, at *Tcrb* and *Igk* loci, loop extrusion appears to proceed in either direction, establishing loops between convergent CBEs that juxtapose V and RAG-bound (D)J RSSs, which allow synapsis by random diffusion and recombination through deletion or inversion^39,41,49^. In this study, we use an integrated functional -omics approach to determine how transcriptional regulatory elements and CBEs at either end of *Tcrb* guide locus folding to facilitate its long-range assembly. The major structural features of folded *Tcrb* loci are: i) homotypic interactions between transcriptionally active Vβ segments and between active Vβ and RC chromatin, ii) loops between convergent Vβ and DJβ CBEs, and iii) segmented stripes of interactions across *Tcrb* loci. The latter feature is consistent with loop extrusion initiating at a Vβ or DJβ CBE and terminating at a DJβ or Vβ CBE, respectively^39^. We demonstrate that the promoter of the most RC-distal Vβ segment, *Trbv1*, functions independent of transcriptional elongation and ncRNA production to recruit NIPBL and cohesion. Promoter-directed loading of NIPBL initiates loop extrusion from the CTCF-bound *Trbv1* 3’CBE to drive *Trbv1* interactions with all other *Tcrb* segments and *Trvb1* rearrangement with DJβ segments. The DJβ CBEs function as terminators or initiaters of loop extrusion and serve as insulators that prevent activation of upstream or downstream promoters by focusing Eβ activity on DJβ promoters. While the DJβ CBEs are dispensable for looping between Vβ segments and the RC, they shape interactions and rearrangements between Vβ and DJβ segments. Finally, we show that the powerful Eβ enhancer element is vital for accumulation of NIPBL and cohesin at the RC and loop extrusion from the RC through Vβ chromatin. However, Eβ is dispensable for forming loops between DJβ and Vβ CBEs, indicating that loop extrusion initiating within Vβ chromatin is sufficient to generate these important architectural features. We conclude that Vβ promoters and Eβ recruit NIPBL and cohesin to initate loop extrusion in opposite directions between Vβ and DJβ chromatin, providing redundant architectural mechanisms for folding *Tcrb* into three dimensional structures that facilitate and shape long-range Vβ-to-RC recombination.

## Results

### Vβ promoter-mediated loading of cohesin drives long-range Vβ-RC interactions

The mouse *Tcrb* locus is well-suited for investigating mechanistic relationships between genome topology and gene regulation for several reasons: i) it is a tractable *in vivo* model for programmed DNA rearrangements that is of manageable complexity, ii) unlike other AgR loci, its RC harbors a single enhancer (Eβ) that mediates all steps of gene assembly, and iii) it harbors two isolated Vβs (*Trbv1* and *Trbv31*) that can be studied without confounding effects of flanking Vβ segments^22,39,50^. *Trbv1* resides 150 kb upstream of the 250 kb Vβ cluster (*Trbv2-30*), which is positioned 250 kb upstream of two DJβ-Cβ clusters (DJβ1-Cβ1 and DJβ2-Cβ1), Eβ, and *Trbv31* (Fig. 1A). In DN thymocytes undergoing active *Tcrb* gene assembly, Eβ contacts Dβ promoters to activate transcription, RAG accessibility, and RC formation at DJβ segments, thereby driving Dβ-to-Jβ rearrangements^33,51–55^. Vβ promoter activation and subsequent rearrangement of the corresponding *Trbv* appears to be Eβ-independent, although the latter relationship has been tested for only a limited number of *Trbv* gene segments^56,57^. Independent of RAG binding,*Tcrb* adopts topologies in DN thymocytes wherein transcriptionally active Vβ and DJβ chromatin regions interact with each other. These active blocks segregate spatially from intervening inactive chromatin spanning two blocks of silent trypsinogen (*Prss*) genes, one situated between the Vβ cluster and *Trbv1*, and a second between the Vβ and DJβ1 clusters^22,39^. *Tcrb* harbors 18 CBEs among the upstream Vβ segments that reside in convergent orientation relative to three CBEs flanking the DJβ clusters: 5’PC and CBE1 residing 27 kb and 3 kb upstream of Dβ1, respectively, and CBE3 between Eβ and *Trbv31* (Fig. 1A)^22,58^. The locus also contains a CBE (CBE2) between CBE1 and Dβ1 that lies in the same orientation as the Vβ CBEs^22,58^. In the DN thymocyte population, cohesin and CTCF bind at each of the 22 *Tcrb* CBEs^58^ and chromosome loops form between numerous Vβ and DJβ CBEs, potentially arising via loop extrusion proceeding across *Tcrb* from V→RC and/or RC→V^39^. In this regard, both the 3’CBE and promoter of *Trbv1* are required for its interactions with other *Tcrb* segments and its recombination with DJβ segments^39^. However, the functions of *Tcrb* promoters, resultant ncRNAs, CBEs, and the Eβ enhancer in regulating topology and recombination of *Tcrb* are largely unknown.

**Figure 1.**
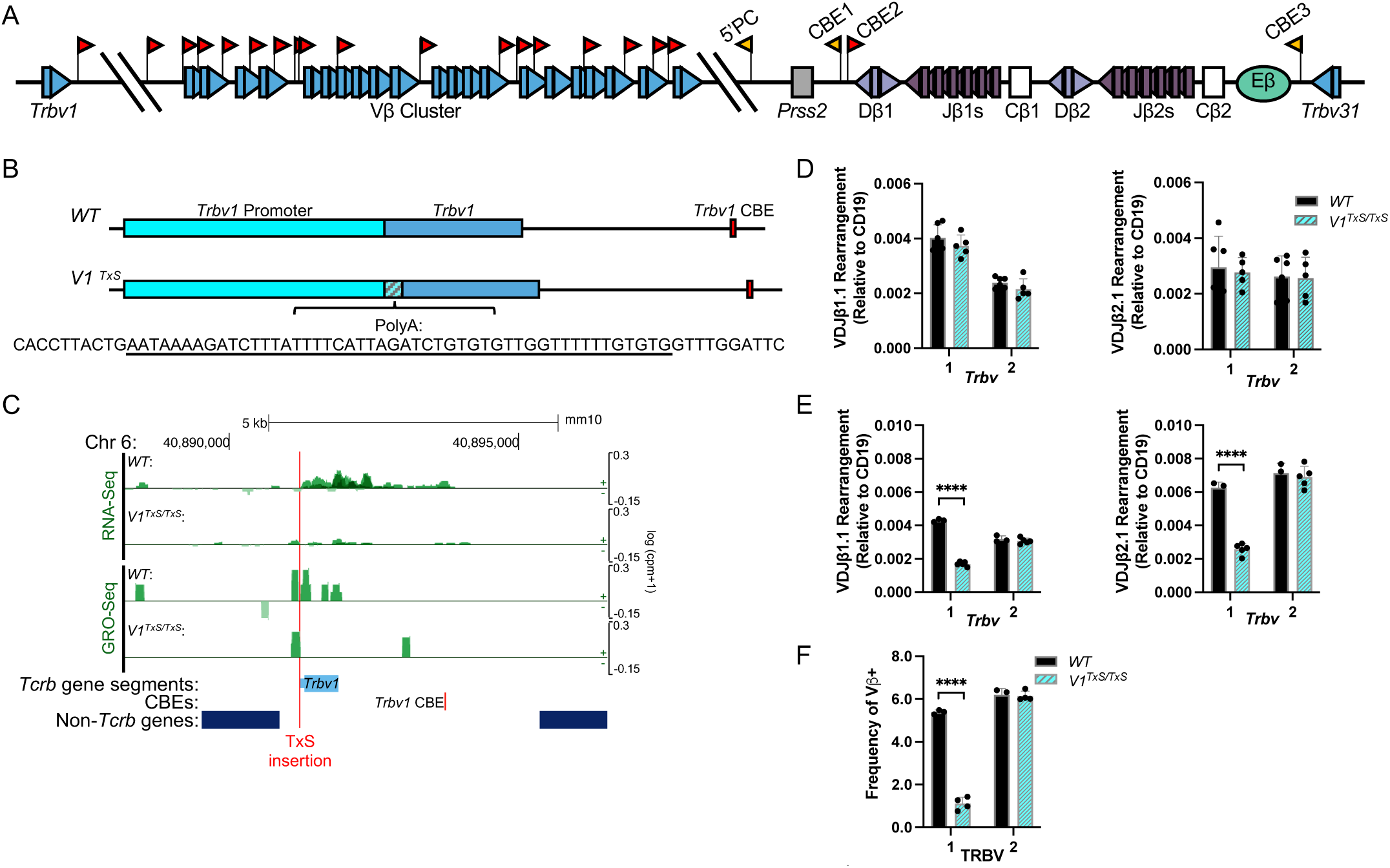
*Trbv1* transcriptional elongation is dispensable for *Trbv1* recombination with DJβ segments. **(A)** Schematic representation of *Tcrb* genomic organization. **(B)** Schematic representation of *Trbv1* on *WT* or *V1^TxS^* alleles. Inserted poly(A) sequence (TxS) is displayed. **(C)** Genome browser tracks of mouse chromosome 6 zoomed to *Trbv1* depicting two replicates of RNA-Seq or GRO-Seq (sense and anti-sense transcripts) for *WT* or *V1^TxS/TxS^* DN cells. The red line indicates TxS placement. **(D)** Taqman qPCR quantification of *Trbv1* and *Trbv2* rearrangements to Jβ1.1 or Jβ2.1 in *WT* or *V1^TxS/TxS^*DN3 thymocytes (n=6,5) or **(E)** total thymocytes (n=3, 5). Multiple unpaired t tests; ****p<0.0001. **(F)** Quantification of TRBV1^+^TCRβ^+^ and TRBV2^+^TCRβ^+^ thymic αβ T cells in *WT* (n=3) or *V1^TxS/TxS^* (n=4) mice. Multiple unpaired t tests; ****p<0.0001.

To elucidate mechanisms by which the *Trbv1* promoter drives recombination, we inserted a transcriptional terminator sequence (TTS) immediately downstream of the *Trbv1* transcriptional start site in the mouse germline (Fig. 1B, *V1^TxS^*allele). We envisioned that the inserted TTS would block transcription of ncRNAs through the *Trbv1* gene segment without otherwise affecting the activation status of *Trbv1* chromatin. We analyzed *V1^TxS/TxS^* mice on a *Rag1^-/-^*background to block thymocyte development at the DN stage and avoid complications of different genomic and topological configurations of *Tcrb* arising from V(D)J recombination. RNA-Seq and Gro-Seq performed on *V1^TxS/TxS^*and *WT* cells validated that the inserted TTS impairs *Trbv1* promoter-mediated transcription through the germline *Trbv1* gene segment, dramatically decreasing the level of *Trbv1* ncRNA (Fig. 1C). Consistent with this, CUT & RUN revealed lower levels of the elongating form of RNA polymerase II (Pol2Ser2) across *Trbv1* in *V1^TxS/TxS^* cells as compared to *WT* cells (Fig. 2A). Despite decreased levels of *Trbv1* transcription in *V1^TxS/TxS^* DN cells, the histone H3K27ac modification associated with transcriptionally active chromatin was relatively unaffected at *Trbv1* (Fig. 2A). MicroCapture C (MCC), a method to visualize frequent or robust interactions between specified genomic regions^59^, revealed strikingly similar patterns of *Tcrb* contacts in *V1^TxS/TxS^* and *WT* DN thymocyes (Fig. 2B). Notably, in both cell types, transcriptionally active *Trbv1* chromatin generated a path of contacts with active chromatin over the Vβ cluster and RC, but interacted less with intervening silent chromatin, forming a segmented stripe originating at *Trbv1* (Fig. 2B). This architectural feature likely forms through cohesin-mediated loop extrusion initiating at the *Trbv1* 3’CBE and extending downstream to DJβ CBEs, with this process impeded more often when translocating across active chromatin or encountering CTCF-occupied CBEs^39^. Consistent with normal interactions between *Trbv1* and DJβ segments, *V1^TxS/TxS^*mice on a RAG-sufficient background showed normal levels of *Trbv1* to-DJβ rearrangements in DN thymocytes (Fig. 1D). We conclude that neither readthrough transcription nor the resultant ncRNAs emanating from the *Trbv1* promoter are required for cohesin-directed loop extrusion from the *Trbv1* 3’CBE or its long-range contacts and recombination with distal DJβ segments. Notably, despite normal levels of *Trbv1* rearrangements in non-selected DN thymocytes, *V1^TxS/TxS^* mice had lower levels of *Trbv1* rearrangements in total thymocytes and a reduced frequency of mature αβ T cells expressing *Trbv1* in their surface AgRs (Fig. 1E, F). This deficit was anticipated because transcription through the *Trbv1* coding sequence is necessary for expression of the rearranged variable region exon into a functional mRNA.

**Figure 2.**
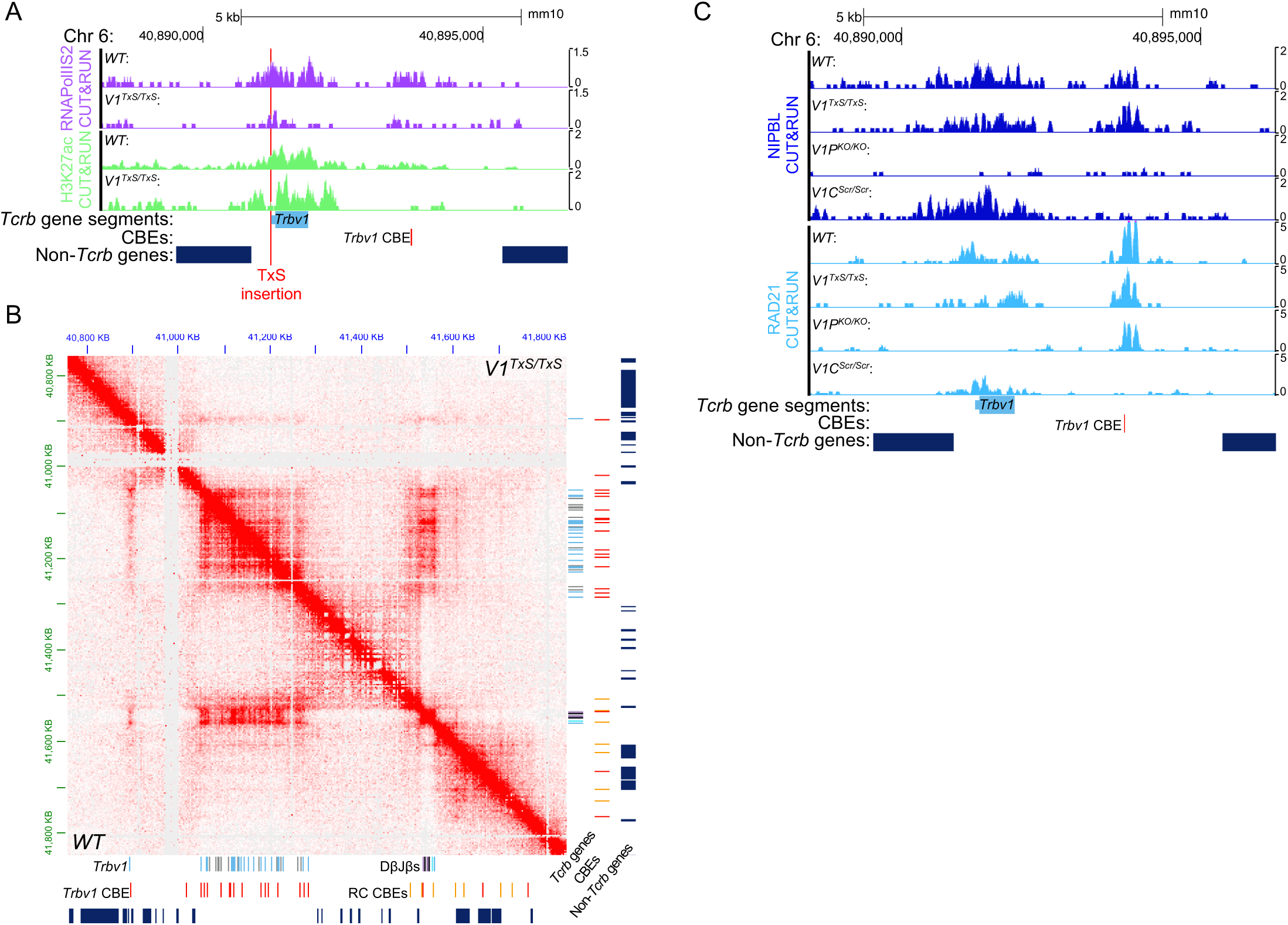
The *Trbv1* promoter loads NIPBL and cohesin to initiate loop extrusion. **(A)** Genome browser tracks of *Trbv1* depicting a representative of two replicates of CUT&RUN for RNAPolIISer2 and H3K27ac in *WT* or *V1^TxS/TxS^* DN cells. **(B)** Micro-Capture C (MCC) heat map of *WT* (bottom, left panel) or *V1^TxS/TxS^* (top, right panel) *Tcrb* loci in DN thymocytes. Data are combined from two independent experiments, each performed on cells pooled from at least five *WT* or *V1^TxS/TxS^* mice of either sex presented at 3 kb resolution. **(C)** Genome browser tracks of *Trbv1* depicting a representative of two replicates of CUT&RUN for NIPBL and RAD21 for *WT, V1^TxS/TxS^, V1P^KO/KO^*, or *V1C^Scr/Scr^* mice.

In contrast to TTS insertion, complete deletion of the *Trbv1* promoter abrogated the segmented stripe of contacts originating from *Trbv1* but had no impact on CTCF binding at the neighboring 3’CBE^39^. Similarly, deletion of the 3’CBE abrogated stripes emanating from *Trvb1* but had no significant effect on transcription of the gene segment. To dissect functions of the *Trbv1* promoter and 3’CBE in supporting loop extrusion, we more deeply characterized the recruitment of architectural factors at *V1^TxS^*, *Trbv1* promoter deleted (*V1P^KO^*), *Trbv1* 3’CBE (*V1C^Scr^*), and wild-type (*WT*) *Tcrb* alleles *in vivo*, with these alleles homozygous on the *Rag1^-/-^* background. We found that the RAD21 subunit of cohesin and the NIPBL cohesin loader accumulate similarly over the promoter, gene segment, and 3’CBE of *Trbv1* in *V1^TxS/TxS^* and *WT* thymocytes, consistent with their retention of segmented stripes originating at *Trbv1* (Fig. 2C). NIPBL binding was lost throughout *Trbv1* chromatin in *V1P^KO/KO^* thymocytes, which also had a corresponding decrease in RAD21 occupancy at the *Trbv1* promoter and gene segment, but retained relatively normal RAD21 binding at the 3’CBE (Fig. 2C). In contrast, for *V1C^Scr/Scr^*cells, we detected relatively normal accumulation of NIPBL and RAD21 at the *Trbv1* promoter and gene segment, but a loss of RAD21 binding at the 3’CBE (Fig. 2C). Together, our new and published data support a mechanism whereby the *Trbv1* promoter directs loop extrusion initiating from the *Trbv1* CBE by loading NIPBL and RAD21 over the *Trbv1* gene segment. This topological function of the *Trbv1* promoter does not require transcriptional elongation but does require CTCF occupancy of the *Trbv1* CBE to serve as an anchor for cohesin-mediated loop extrusion.

### The DJβ CBEs modulate Vβ and Dβ-Jβ gene segment usage during *Tcrb* gene assembly

Notable architectural features of *Tcrb* loci in DN thymocytes include trails of interactions originating in Vβ or DJβ CBEs, and passing through transcriptionally active DJβ or Vβ chromatin, respectively, but these trails are interrupted by intervening silent chromatin regions (Fig. 2B)^39^. Together, these interactions form segmented stripes that include chromosome loops between convergent Vβ and DJβ CBEs. As shown for the *Trbv1* 3’CBE^39^, these structures likely reflect cohesin-mediated loop extrusion initiating from a Vβ or DJβ CBE and progressing downstream or upstream, respectively, with less impeded progress when transversing silent chromatin. We envision that loop extrusion proceeds in one direction or the other within an individual cell, leading to a pattern of bidirectional loop extrusion across the entire DN thymocyte population. In this regard, the 5’PC, CBE1, and CBE3 DJβ CBEs should function as initiators and/or terminators of loop extrusion to establish loops between Vβ and DJβ chromatin, facilitating long-range Vβ rearrangements.

To test this model, we generated and analyzed mice with deletion of 5’PC, CBE1, or CBE3 individually or in combination (Fig 3A). We first used Adaptive Immunosequencing, a commercial Next-Generation Sequencing (NGS) platform, on DNA isolated from non-selected DN thymocytes to determine the effect of deleting DJβ CBEs on gene segment utilization in *Tcrb* rearrangements. Mice with homozygous deletion of 5’PC (*5’PC^KO/KO^*) or CBE1 (*C1^KO/KO^*) exhibited a moderate decrease in usage of the most RC-distal Vβs (*Trbv1-Trbv3*) and a corresponding increase in usage of the most RC-proximal Vβ cluster segments (*Trbv26-Trbv30*) (Fig 3B). In contrast, mice with homozygous deletion of CBE3 (*C3^KO/KO^*) had a ∼2-fold increase in *Trbv31* usage and no discernable effect on rearrangement of other Vβ segments (Fig. 3B). Notably, only *C1^KO/KO^* mice exhibited decreased usage of DJβ1 versus DJβ2 gene segments (Fig. 3C). Mice with homozygous deletion of 5’PC, CBE1, and CBE3 (*513^TKO/TKO^*) showed dramatically increased usage of RC-proximal Vβ cluster segments, most notably of *Trbv30* as compared to *WT*, *5’PC^KO/KO^*, or *C1^KO/KO^*mice, but normal utilization of *Trv31* (Fig. 3B). These data indicate that 5’PC and CBE1 exert unique and redundant functions in controlling rearrangements of upstream Vβ segments, whereas CBE1 and CBE3 have a unique function in regulating recombination of DJβ or *Trbv31* segments, respectively.

**Figure 3.**
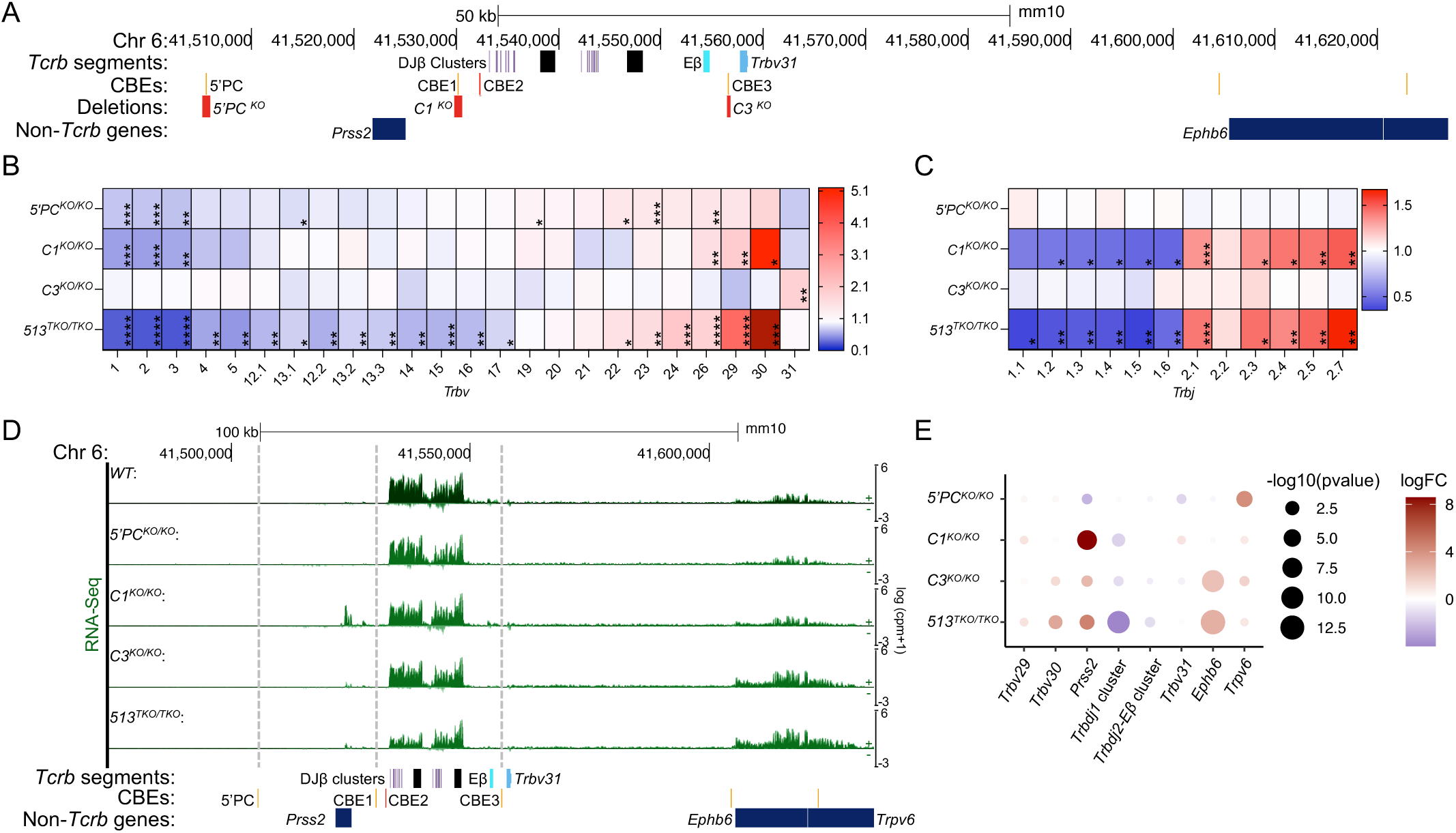
DJβ CBEs promote a broad Vβ repertoire and restrict Eβ activity. **(A)** Genome Browser view of mouse chromosome 6 spanning the *Tcrb* DJβ gene segments with tracks presenting nucleotide position, *Tcrb* gene segments (Dβ segments, lavender; Jβ segments, purple; Cβ exons, black; Eβ, cyan; *Trbv31* segment, light blue), CBEs (red for sense orientation or orange for antisense orientation), the deleted regions in the indicated CBE mutant (red) and non-*Tcrb* genes. **(B)** Quantification of the percentage of total unique *Tcrb* genes involving each Vβ or **(C)** Jβ gene segment, baseline-corrected to *WT* from Adaptive Immunosequencing performed on DNA isolated from DN thymocytes of *WT*, *5’PC^KO/KO^*, *C1^KO/KO^*, *C3^KO/KO^*, or *513^TKO/TKO^* mice. n=2, multiple unpaired t-tests. *p<0.05, **p<0.01, ***p<0.001, ****p<0.0001. Dark red is off scale; *Trbv30* value in *513^TKO/TKO^* mice is 20.9. **(D)** Genome browser tracks of the *Tcrb* DJβ region depicting two replicates each of RNA-Seq (sense and anti-sense transcripts) for *WT*, *5’PC^KO/KO^*, *C1^KO/KO^*, *C3^KO/KO^*, or *513^TKO/TKO^* mice. **(E)** Quantification of transcription for DJβ gene segments and adjacent genes, logFC over *WT*.

### CBE1 and CBE3 restrict activity of the Eβ enhancer to Dβ promoters

To determine how the DJβ CBEs regulate *Tcrb* rearrangements, we generated and analyzed *5’PC^KO/KO^*, *C1^KO/KO^*, *C3^KO/KO^*, and *513^TKO/TKO^* mice on the *Rag1^-/-^* background. Due to numerous correlations between rearrangement and germline transcription of gene segments^60^, we conducted RNA-Seq on DN thymocytes of these strains to monitor *Tcrb* transcripts. While the *5’PC^KO/KO^* and *C3^KO/KO^* strains displayed normal levels of transcripts from *Tcrb* segments, *C1^KO/KO^* mice exhibied lower levels of DJβ1 transcripts (Fig. 3D, E). The *C1^KO/KO^* mice also exhibited increased transcripts of the *Prss2* gene located upstream of CBE1, whereas *C3^KO/KO^* mice displayed more transcripts of the *Ephb6* gene residing downstream of *Trbv31* (Fig. 3D, E). The *513^TKO/TKO^* mice exhibited more accentuated effects on DJβ1, *Prss2*, and *Ephb6* transcripts, as well as slightly lower levels of DJβ2 transcripts (Fig. 3D, E). These data demonstrate that CBE1 and CBE3 serve as insulators that protect adjacent non-*Tcrb* genes from Eβ function, while CBE1 also focuses Eβ activity to the Dβ1 promoter.

### The DJβ CBEs serve as chromosome loop anchors that establish diverse *Tcrb* topologies

To determine how the DJβ CBEs participate in the folding of *Tcrb* loci, we analyzed *Tcrb* topology by MCC and binding of chromosome looping factors by CUT&RUN in DN thymocytes from RAG-deficient *WT* or *513^TKO/TKO^* mice. In the *WT* cells, we found that CTCF, RAD21, and NIPBL bind throughout *Tcrb* mainly at sequences harboring CBEs (Fig. 4A). RAD21 and NIPBL also bind over Eβ and transcribed DJβ and *Trbv31* segments in *WT* cells (Fig. 4A). In *513^TKO/TKO^* cells, *Tcrb* loses CTCF, RAD21, and NIPBL binding over specific sites (Fig. 4A). Neither CTCF nor RAD21 accumulate over regions spanning the deleted 5’PC, CBE1, or CBE3 elements. Moreover, neither RAD21 nor NIPBL bind to DJβ1 segments that are transcribed at lower-than-normal levels in *513^TKO/TKO^* cells (Fig. 4A; 3D,E). <u>Notably, CTCF, RAD21 and NIPBL were all bound at largely normal levels across the Vβ segments, with only RAD21 bound at a higher level near both *Trbv29* and *Trbv30,* correlating with their increased rearrangement (Fig. S1A).</u> MCC data revealed altered *Tcrb* topologies in *513^TKO/TKO^* cells. The most striking topological differences between *WT* and *513^TKO/TKO^* DN thymocytes were that the latter lacked interaction trails originating at the deleted CBE regions and traversing through Vβ chromatin (Fig. 4B, black arrows). Instead, we observed a broader trail of contacts between sequences spanning the DJβ2 –Eβ region and Vβ chromatin in *513^TKO/TKO^* cells (Fig. 4B). These obsevations are supported by virtual 4C views, wherein interactions between the Vβ cluster and DJβ1, but not the DJβ2 segments, are lower within *513^TKO/TKO^* cells as compared to *WT* cells (Fig. 4C). While *513^TKO/TKO^* DN cells contained no loops between Vβ CBEs and sequences spanning the deleted DJβ CBEs, they did maintain loops between Vβ CBEs and sequences spanning DJβ2-Eβ (Fig. 4C,D). Another notable difference for *513^TKO/TKO^* loci was an increase in interaction, including loops, between Vβ chromatin and sequences downstream of *Tcrb*, especially the convergent CBE of the active *Ephb6* gene, in *513^TKO/TKO^*cells (Fig. 4C,D), which is consistent with loop extrusion intiating at Vβ CBEs extending more often past *Tcrb*. Finally, consistent with the above elucidated roles of CBE1 and CBE3 as insulators, we observe sharp segregation between DJβ segments and their flanking chromatin regions in *WT* but not *513^TKO/TKO^*cells (Fig. 4B, blue brackets). Zooming into the 5’PC-CBE3 region (Fig. 4D), we found that Eβ loops to both the DJβ1 and DJβ2 segments in *WT* cells, but only to the DJβ2 segments in *513^TKO/TKO^* cells, suggesting that these CBEs play a role in directing Eβ activity. Collectively, these MCC data support our model wherein DJβ CBEs function as both insulators of Eβ that delineate the active DJβ segments from the flanking silent chromatin, as well as initiators and/or terminators of loop extrusion to establish loops between Vβ and DJβ chromatin.

**Figure 4.**
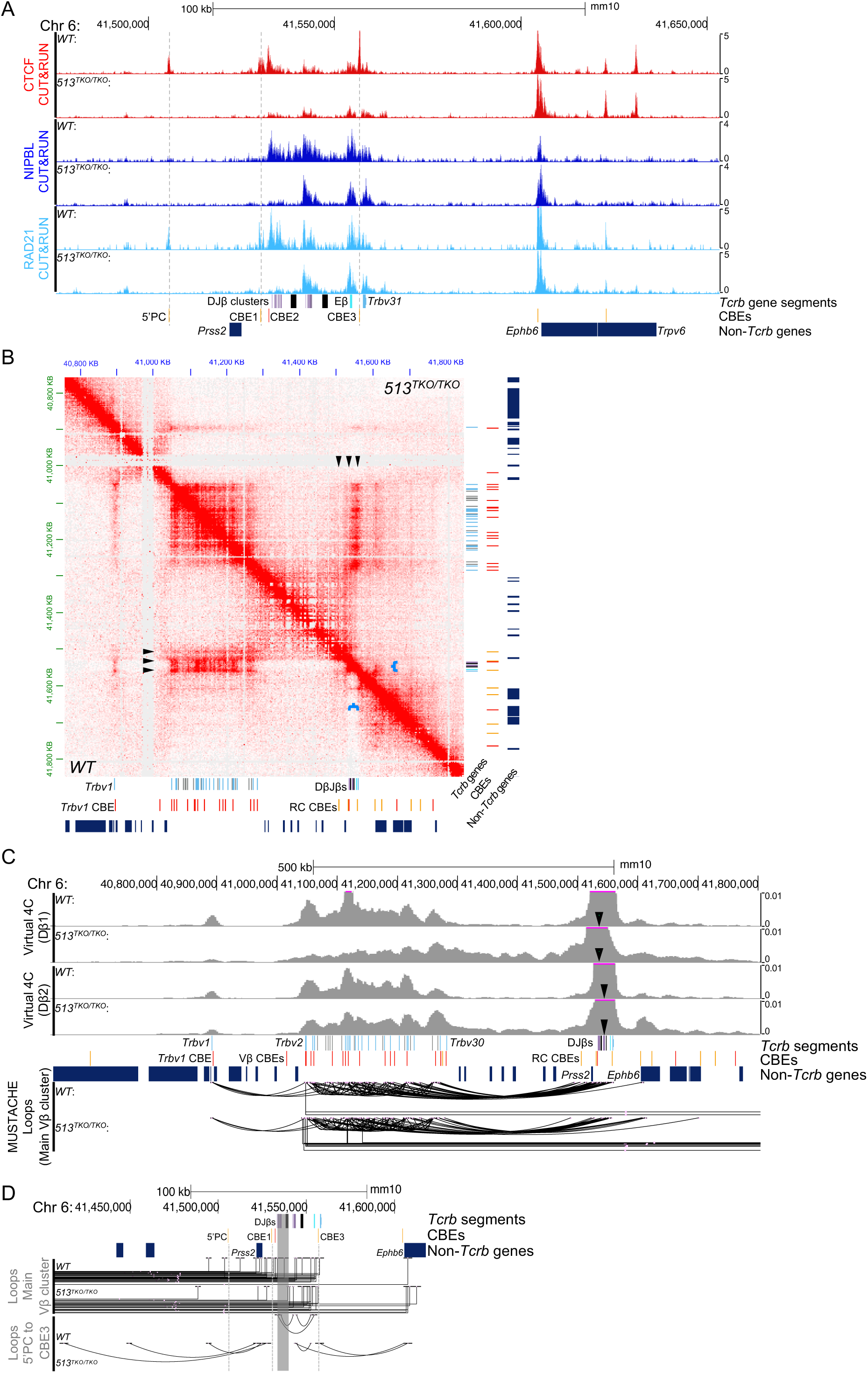
DJβ CBEs focus cohesin loading and anchor Vβ-DJβ loops. **(A)** Genome browser tracks of *Tcrb,* zoomed in to the DJβ segments, depicting a representative of two replicates of CUT&RUN for CTCF, NIPBL or RAD21 for *WT* or *513^TKO/TKO^* mice. **(B)** MCC heat map of *WT* (bottom, left panel) or *513^TKO/TKO^* (top, right panel) *Tcrb* loci in DN thymocytes. Data are combined from two independent experiments, each performed on cells pooled from at least five *WT* or *513^TKO/TKO^* mice of either sex presented at 3 kb resolution. The positions of the DJβ CBEs are indicated (black arrows), as well as the discussed segregation of the DJβ clusters from the flanking silent regions (blue brackets). **(C)** Genome browser tracks depicting virtual 4C tracks from the Dβ1 or Dβ2 viewpoint from *WT* or *513^TKO/TKO^* thymocytes generated from MCC data in panel B (black arrows indicate the 4C viewpoint). Below the gene tracks, MUSTACHE loops are represented as arcs from MCC data from panel B, filtered by interactome so that least one anchor falls within the main Vβ cluster. **(D)** Genome browser tracks of *Tcrb,* zoomed in to the DJβ segments, depicting MUSTACHE loops from MCC data from panel B, filtered by interactome where at that least one loop anchor falls within the main Vβ cluster (top tracks) or between 5’PC and CBE3, with both anchors falling in the zoomed in region (bottom tracks).

### Eβ is necessary for loop extrusion initiating from DJβ chromatin but dispensable for *Tcrb* folding

Enhancer-mediated activation of (D)J chromatin is pivotal for AgR gene assembly by enabling RAG binding of (D)J RSSs to form the RC^33^. However, the roles of (D)J-proximal enhancer elements in facilitating the formation of long-range contacts within AgR loci remain enigmatic due to several complicating factors. These include the presence of multiple enhancers in most AgR loci and possible distinctions between locus contraction, as measured by FISH, and precise V-RC contacts^22,30,34,36–38^. Among AgR loci, *Tcrb* is unique in that it has only one (D)J-proximal enhancer, Eβ, which singularly activates RC chromatin and all steps of *Tcrb* gene assembly^61,62^. While Eβ is dispensable for point-to-point contacts between several Vβ segments and the RC^22^, its mechanistic role in loop extrusion and the formation of *Tcrb* topologies remains unknown.

To address this knowledge gap, we generated *Eβ^KO/KO^*mice on a *Rag1^-/-^* background, hereafter referred to as *Eβ^KO/KO^* mice^61,63^ (Fig. 5A). As shown in Fig. 5B, *Eβ^KO/KO^* DN thymocytes displayed a nearly complete loss of both DJβ transcripts and H3K27ac across the RC, confirming prior studies showing a dominant role for this enhancer in activating Dβ promoters and DJβ chromatin (Fig. 5B). Also, consistent with published findings^22^, deletion of Eβ had no obvious effect on levels of transcripts from Vβ segments, except *Trbv31*, which is located only 3 kb downstream of Eβ (Fig. 5B). To determine if, like the promoter of *Trbv1,* Eβ driven transcriptional activity directs the loading of cohesin by NIPBL at the DJβ region, we performed CUT and RUN for these factors in the *Eβ^KO/KO^* mice. In WT thymocytes, NIPBL and RAD21 were bound at high levels across the RC, including the DJβ segments, Eβ, and *Trbv31* consistent with the current model that NIPBL loads cohesin at regions of transcriptional activity^9^. However, unlike our findings at *Trbv1* where NIPBL was bound to both the *Trbv1* gene segment and adjacent CBE in *WT* cells, the DJβ CBEs were bound by RAD21, but not NIPBL (Fig. 5B). In *Eβ^KO/KO^* cells, CTCF and RAD21 bound at lower levels to 5’PC, CBE1, and CBE3, but, strikingly, neither RAD21 nor NIPBL bound to DJβ segments and Eβ (Fig. 5B). Together, these data support a mechanism whereby Eβ activity directs loading of cohesin by NIPBL at the DJβ end of *Tcrb,* which could initate loop extrusion to extend into the Vβ segments, to generate short-range loops between Dβ and Jβ segments, or both. Importantly, we observed no differences in the binding of CTCF, RAD21, or NIPBL throughout Vβ chromatin between *WT* and *Eβ^KO/KO^* DN thymocytes (Fig. S1B). Thus, in the absence of Eβ activity, any contacts between Vβ and DJβ segments are likely the result of Vβ promoter-directed loading of cohesin and resulting initiation of loop extrusion from Vβ CBEs through DJβ CBEs.

**Figure 5.**
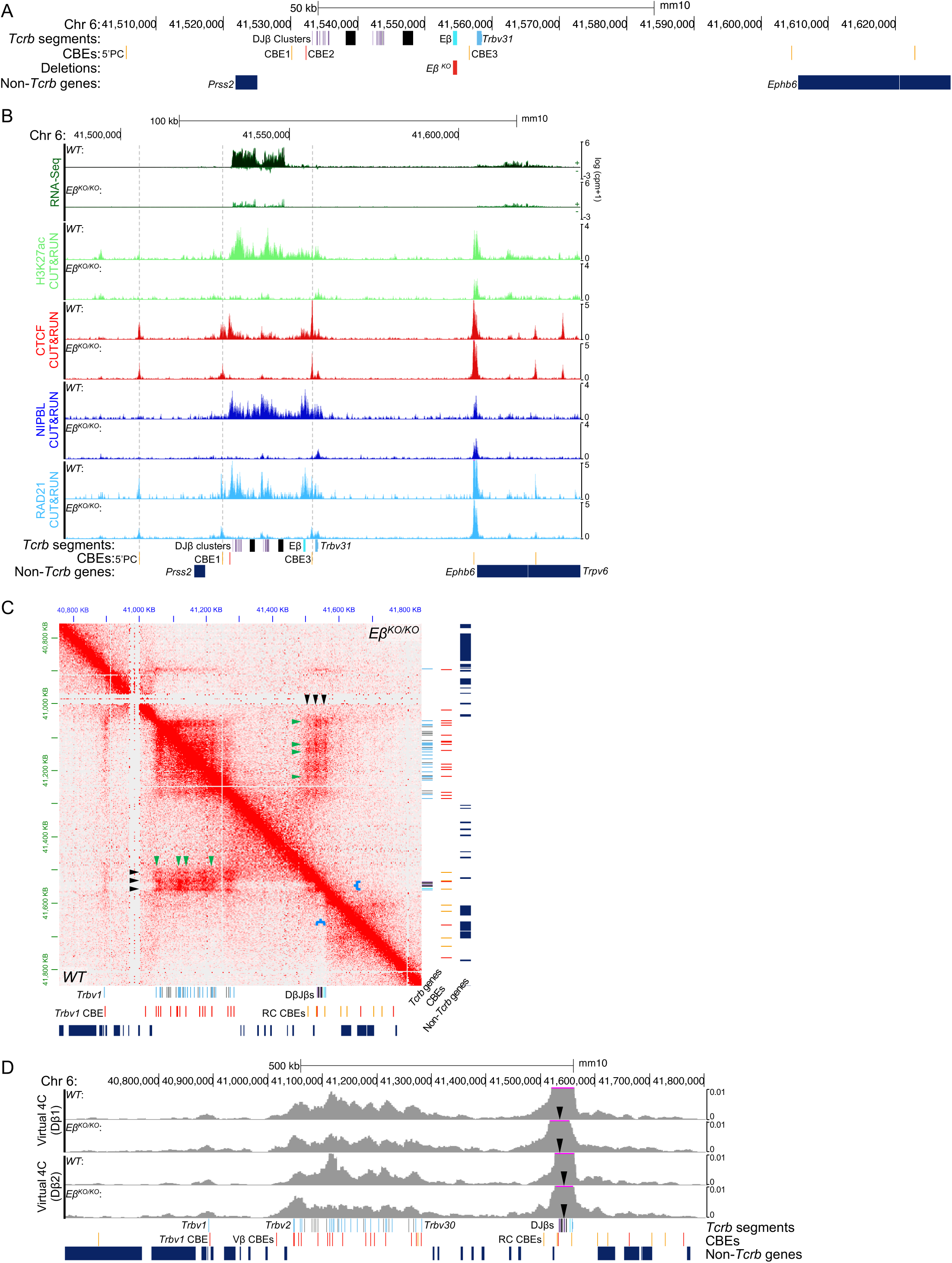
Eβ activation of DJβ clusters directs cohesin loading but is largely dispensable for DJβ CBE loops. **(A)** Genome Browser view of mouse chromosome 6 spanning the *Tcrb* DJβ gene segments with tracks presenting nucleotide position, *Tcrb* gene segments (Dβ segments, lavender; Jβ segments, purple; Cβ exons, black; Eβ, cyan; *Trbv31* segment, light blue), CBEs (red for sense orientation or orange for antisense orientation), the deleted regions in the indicated CBE mutant (red) and non-*Tcrb* genes. **(B)** Genome browser tracks of *Tcrb,* zoomed in to the DJβ segments, depicting two replicates each of RNA-Seq (sense and anti-sense transcripts) or a representative of two replicates of CUT&RUN for H3K27ac, CTCF, NIPBL, or RAD21 for *WT* or *Eβ^KO/KO^* mice. **(C)** HiC heat maps of *WT* (bottom, left panel) or *Eβ^KO/KO^* (top, right panel) *Tcrb* loci in DN thymocytes. Data are combined from two independent experiments, each performed on cells pooled from at least five *WT* or *Eβ^KO/KO^*mice of either sex presented at 3 kb resolution. The positions of the CBEs are indicated (black arrows), as well as the discussed interactions between the DJβs and transcriptionally active Vβs (green arrows) and segregation of the DJβ clusters from the flanking silent regions (blue brackets). **(D)** Genome browser tracks depicting virtual 4C tracks from the Dβ1 or Dβ2 viewpoint from *WT* or *Eβ^KO/KO^*thymocytes generated from HiC data in panel C (black arrows indicate the 4C viewpoint).

In this regard, *Tcrb* topology was largely unchanged by the Eβ deletion when compared with *WT* thymocytes (Fig. 5C). However, there were two notable differences. First, similar to the *513^TKO/TKO^* cells, the DJβ region did not sharply segregate from the flanking, inactive sequences in the *Eβ^KO/KO^* cells (Fig. 5C, blue brackets). As described above, this loss in the *513^TKO/TKO^* cells suggested that the CBEs aided in delineating the active DJβ chromatin from the surrounding inactive sequences; whereas, in the *Eβ^KO/KO^* cells, the DJβ chromatin is no longer active and thus, doesn’t segregate from the neighboring inactive regions (Fig. 5C, blue brackets). The second difference was that the trail of contacts initiating at CBE3 and extending to 5’PC in *WT* cells was absent in *Eβ^KO/KO^*thymocytes. This latter finding is consistent with a necessity for cohesin loading at the DJβ segments to initiate loop extrusion and generate short range loops among DJβ segments, or to extend extrusion through the Vβ segments, or both (Fig. 5C). However, despite the loss of cohesin loading at the DJβ region, *Eβ^KO/KO^*cells retained loops between convergent Vβ and DJβ CBEs and the three stripes corresponding to interactions between the DJβ CBEs and the Vβ chromatin (Fig. 5C, black arrows). This observation indicates that the contact stripes are not formed from loop extrusion initiating at the DJβ CBEs, but instead from loop extrusion initiating at Vβ chromatin and terminating at these CBEs. Finally, there are less robust interactions between the highly transcribed Vβ segments and the DJβ chromatin in the *Eβ^KO/KO^* (Fig 5C, green arrows) suggesting homotypic interactions, while not necessary for long-range interactions between Vβ and DJβ segments, can help bolster contacts between active Vβ and DJβ chromatin. Altogether, these data indicate that independent of Eβ function, Vβ promoter-directed loading of cohesin and resulting initiation of loop extrusion from Vβ CBEs through DJβ CBEs is sufficient to juxtapose Vβ and DJβ segments for rearrangement.

## Discussion

The dynamic reorganization of chromosome topology regulates many biological processes, including lymphocyte AgR gene assembly through RAG-mediated recombination of gene segments, which can be separated by vast linear genomic distances. Loop extrusion and homotypic chromatin interactions have each been implicated in driving the long-range *Tcrb* interactions that are required for recombination between accessible Vβ segments and RAG-bound DJβ segments^39^. In this study, we integrated genetic and -omics approaches to elucidate how a Vβ promoter, the DJβ CBEs, and the Eβ enhancer orchestrate intra-*Tcrb* interactions and thereby shape long-range *Tcrb* recombination. We discovered that the *Trbv1* promoter functions independently of transcriptional elongation to load cohesin and, consequently, orchestrates chromosome loop extrusion initiating at the *Trbv1* 3’CBE and progressing downstream to convergent DJβ CBEs. The latter modulate *Tcrb* repertoires by serving as terminators of loop extrusion originating from upstream Vβ CBEs, placing these gene segments into spatial proximity of the RC; but the DJβ CBEs also serve as insulators that focus Eβ activity to DJβ promoters. We find that Eβ, which activates DJβ chromatin in DN thymocytes to enable RC formation, is essential for binding of NIPBL and cohesin over its composite gene segments, a prerequisite for loop extrusion to initiate within the RC. Surprisingly, however, the Eβ super-enhancer is dispensable for generating long-range Vβ to RC contacts. Thus, chromosome looping originating in the Vβ cluster and proceeding downstream to the RC is sufficient to mediate normal long-range contacts between Vβ and DJβ segments independent of homotypic interactions nucleated by Eβ-dependent activation of RC chromatin. Collectively, our study demonstrates that Vβ promoter-mediated cohesin loading and resultant loop extrusion from an adjacent Vβ CBE cooperate with DJβ CBEs, which function as terminators of loop extrusion and chromatin insulators, to establish *Tcrb* contacts that facilitate assembly of a broad *Tcrb* gene repertoire.

We employed the most distal *Trbv1* gene segment as a stand-alone model to resolve mechanisms of *Tcrb* folding and recombination. We found that its associated promoter directs loop extrusion initiating from the *Trbv1* 3’CBE by loading NIPBL and cohesin, a process that is independent of transcriptional elongation. This topological function of the *Trbv1* promoter requires CTCF binding at its adjacent CBE, which serves as an anchor for cohesin-mediated loop extrusion progressing to DJβ CBEs, thereby forming chromosome loops that juxtapose *Trbv1* with the distal RC. In embryonic stem cells, promoter and enhancer activity has been implicated in transcription-dependent binding of NIPBL^9^ and cohesin loading, sprouting models wherein active promoters load cohesin onto chromosomes and/or transcription elongation pushes unidirectional cohesin migration during loop extrusion^16,17,64,65^. However, our data indicate that transcriptional elongation through *Trbv1* is dispensible for cohesin loading by the *Trbv1* promoter, its migration downstream to the *Trbv1* 3’CBE, and subsequent loop extrusion. *In vitro,* NIPBL is necessary for cohesin-mediated loop extrusion^10^, implying that the *in vivo* process originates from CBEs bound by CTCF, NIBPL, and cohesin. This further implies that CBEs occupied by CTCF and cohesin without NIPBL, can function as terminal, but not initiating, anchors for loop extrusion. In this context, the persistence of RAD21 at the *Trbv1* 3’CBE even when the *Trbv1* promoter is deleted has two potential explanations: i) cohesin loaded elsewhere translocates to this CBE but cannot initiate loop extrusion without NIPBL, and/or ii) cohesin-directed loop extrusion initiating elsewhere (e.g., the RC) terminates at this CBE. As we did not find chromosome loops between the *Trbv1* CBE and any downstream *Tcrb* CBEs in *Trbv1* promoter knockouts^39^, the second scenario either is eliminated, or it does not result in stable chromosome loops, perhaps due to lack of homotypic interactions between the now silent *Trbv1* and active RC chromatin domains. Notwithstanding, our data provide strong *in vivo* evidence that cohesin initiates loop extrusion from the *Trbv1* 3’CBE, a process dependent on *Trbv1* promoter-mediated recruitment of NIPBL but independent of transcriptional elongation.

Genetic manipulation of the DJβ CBEs revealed their dual functions as insulators that focus Eβ activity to DJβ promoters, and as terminators of loop extrusion from upstream convergent Vβ CBEs. Each of these functions contributes different aspects to diversification of the final *Tcrb* repertoire, as well as transcriptional regulation of *Tcrb* and genes neighboring the locus. Our data indicate that the DJβ CBEs are important for directing Eβ activity to Dβ promoters, likely by forming stable promoter-enhancer contacts. Accordingly, loss of CTCF binding at CBE1 or CBE3 results in a significant increase in the transcription of genes upstream and downstream, respectively. Moreover, loss of CTCF binding at CBE1 eliminated Eβ to Dβ1 looping, which significantly reduced transcription of the DJβ1 cluster, implying that CBE1 specifically restricts Eβ activity to the Dβ1 promoter. Notably, these transcriptional changes led to differences in *Tcrb* topology and gene segment utilization during long-range Vβ rearrangement. The reduction in DJβ1 transcription corresponded with decreased interactions and rearrangements of DJβ1 and upstream Vβ segments. Furthemore, the loss of CBE1 and 5’PC insulator functions increased transcription, RC interactions, and rearrangement of the most proximal Vβ segments. Finally, the sharp segregation of active RC chromatin in WT thymocytes from upstream silent chromatin was diminished when DJβ CBEs were inactivated, suggesting that these CBEs indirectly shape homotypic interactions by insulating Eβ to partition active and inactive chromatin regions. Moreover, we found that the DJβ CBEs serve as terminators for loop extrusion initiating from upstream Vβ chromatin, and potentially function as initiating anchors for loop extrusion from the RC towards Vβ segments. However, 5’PC and CBE1 are not bound by NIPBL, suggesting that only CBE3 acts as an initiating anchor for cohesin-mediated RC-Vβ loop extrusion. Notably, when all DJβ CBEs are deleted, some Vβ-RC contacts are maintained, but are focused on the DJβ2-Eβ region, which is the most transcriptionally active cluster. The revised contacts are consistent with three potential scenarios: (i) loop extrusion initiating in Vβ chromatin is terminated by active transcription within the DJβ2-Eβ cluster or its homotypic interactions with active Vβ segments, (ii) transcription at DJβ2-Eβ anchors the initiation of loop extrusion that extends to Vβ chromatin in the absence of DJβ CBEs, or (iii) homotypic interactions, drive contacts between Vβ and Dβ-Jβ segments independent of chromosome looping. Consistent with the first scenario, in the absence of the DJβ CBEs, the Vβ cluster interacts more frequently with the convergent CBEs downstream, perhaps because Vβ-directed loop extrusion proceeds further downstream when the RC-flanking CTCF boundaries are removed.

In DN thymocytes, Eβ acts as a super-enhancer, forming a region of highly active chromatin over the entire RC region. We now show that Eβ also is essential for RC association with the cohesin loader NIPBL. The absence of NIPBL and cohesin at the RC in *Tcrb* loci lacking Eβ likely precludes initiation of loop extrusion from the DJβ CBEs upstream to the Vβ cluster, which is supported by an absence of stripes emanating from the RC in Hi-C data. However, in the absence of Eβ, DJβ CBEs retain CTCF and some RAD21, which anchor loops with the Vβ segments. This scenario is consistent with loop extrusion initiating at Vβ CBEs, as confirmed at *Trbv1*, proceeding downstream, and terminating at the DJβ CBEs. Importantly, we observe a comparable number of Vβ/DJβ CBE loops in *WT* and *Eβ^KO/KO^* loci, indicating that loop extrusion originating within the Vβ segments is sufficient for juxtaposing Vβ segments with the RC. This model is supported by our prior studies showing that inactivation of Eβ has little to no impact on wholescale contraction of *Tcrb* in DN thymocytes as evidenced by *in situ* hybridization^22^. By comparison, enhancers located in V chromatin of *Igh* or *Igk* loci drive contact of flanking V segments with downstream D-J segments by blocking loop extrusion originating in the RC^26,27,66^. In contrast, while Eβ is required for activation of DJβ segments, as well as the subsequent separation from the flanking silent regions and initiation of loop extrusion within the RC, it is not required for RC contacts with Vβ segments. Instead, it appears that loop extrusion initiating within the Vβ region can be blocked by either the DJβ CBEs or Eβ-driven transcription.

Our study highlights other important contrasts in folding mechanisms employed by *Tcrb* versus other AgR loci. The *Igh* locus folds predominantly by unidirectional, asymmetrical loop extrusion initiating near the RC, enabling RAG scanning of V_H_ segments as they spool past RAG-bound DJ_H_ complexes^19,46^. In contrast, *Igk* folds by juxtaposed loops of Vκ and Jκ segments that undergo diffusion, as opposed to RAG scanning, synapsis^41,49^. This study, combined with our prior work, indicates that a predominant mechanism by which *Tcrb* folds involves loop extrusion initiating from Vβ CBEs and terminating at DJβ CBEs, a process that is incompatible with RAG scanning. Given this finding, we propose a model in which the specific Vβ segment selected for rearrangement in a given cell depends on where cohesin was loaded. Stochastic mechanisms, such as transcriptional bursting, might direct cohesin loading to an active Vβ segment, initiating loop extrusion from an adjacent CBE and increasing the likelihood for synapsis of neighboring Vβ segments by diffusion. Such a model may also provide an important mechanism for promoting the rearrangement of more distal Vβ segments, which are highly transcribed compared with the proximal segments in the main Vβ cluster, limiting initiation of loop extrusion and over-usage of these proximal segments. Consistent with this, when DJβ CBEs are deleted, Eβ activation extends up to at least *Trbv29,* which coincides with higher levels of its recombination, perhaps due to more frequent loading of cohesin.

Our findings also have implications for cohesin-mediated chromosome looping in general. The *Trvb1* promoter mutants provide support for models in which active transcriptional regulatory elements also function as sites for cohesin loading; however, our data are inconsistent with models invoking transcriptional elongation in propelling loop extrusion. In addition, a role for active transcription in pausing of loop extrusion is supported by the retention of long-range contacts in alleles lacking the usual termination sites (i.e., DJβ CBEs), where contacts are refocused to the more transcriptionally active DJβ 2 cluster. Thus, AgR loci will continue to provide insights into mechanisms that generate and maintain genome topologies, especially with regard to developmental switches from active to inactive structures.

## Materials and Methods

### Mice

Mice used for all experiments were 4-6 weeks old, of mixed sex, and housed under specific pathogen-free conditions at the Children’s Hospital of Philadelphia (CHOP) or the Ohio State University (OSU) College of Medicine. All animal husbandry, breeding, and experiments were performed in accordance with national guidelines and regulations and approved by the CHOP Institutional Animal Care and Use Committee and the OSU Institutional Animal Care and Use Committee. Wild-type (C57BL/6J) and *Rag1^-/-^*(B6.129S7-*Rag1^tm1Mom^*/J) mice were obtained from Jackson Laboratories. The *V1P^KO/KO^* and *V1C^Scr/Scr^* mice were previously described^39^ as well as the *Eβ^KO/KO^* mice^61,63^.We used the Easi-CRISPR method ^67^ of CRISPR/Cas9-mediated genomic editing in C57BL/6 zygotes to create mice with an insertion of the synthetic poly(A) sequence (*V1^TxS^*) or a deletion of the three DJβ CBEs (*5’PC^KO^, C1^KO^,* and *C3^KO^*) plus combined deletion (*513^TKO^)*.

For the *V1^TxS/TxS^* mice, we used CRISPOR to identify a suitable guide sequence on the antisense strand to position the cut site at the promoter-*Trbv1* junction. The crRNAs were purchased from Integrated DNA Technologies (IDT) with the identified guide sequence (see Table 1 for a complete list of oligos). The Easi-CRISPR method uses ssDNA as donors (ssODN) for homology directed repair, which carried a verified synthetic poly(A) sequence^68^. The CHOP Transgenic Core electroporated zygotes with a mixture of ctRNA (8 μM; crRNA +tracrRNA), Cas9 protein (3.2 μM), ssODN (20 μM) in Duplex Buffer diluted 1:1 with Opti-MEM buffer. A similar approach was used to generate the knockout mice, with the exception that two different guide RNAs were used to generate a deletion between the two cut sites, with no homology directed repair. The mixture used to electroporate zygotes contained two different ctRNAs (4 μM each; two different crRNAs independently annealed to tracrRNA) and no ssODN. *V1^TxS/TxS^* founders were screened by PCR of tail DNA with the V1TxS5’ and TxSRev or TxSFwd and V1PKO3’ primers to test for the presence of the synthetic poly(A) sequence and to check for appropriate insertion of the 5’ and 3’ homology arms, respectively. The individual or combined deletions of 5’PC, CBE1, and CBE3 were screened with 5’PCKO5’ and 5’PCKO3’, C1KO5’ and C1KO3’, or C3KO5’ and C3KO3’, respectively. For each mutant, two founder lines were selected and backcrossed to *WT* mice for two generations to ensure the propagation of a single validated mutant allele and limit potential off-target CRISPR effects. We crossed heterozygous F2 mice to generate homozygous mice, and genotypes were verified by PCR with the flanking primers, followed by Sanger Sequencing from either end of the PCR product. The sequence-validated founder lines were analyzed by flow cytometry, using *WT* littermate controls, to confirm the same phenotype between founder lines. One line was selected for Adaptive Immunosequencing analysis, and to backcross onto the RAG-deficient background.

**Table 1.**
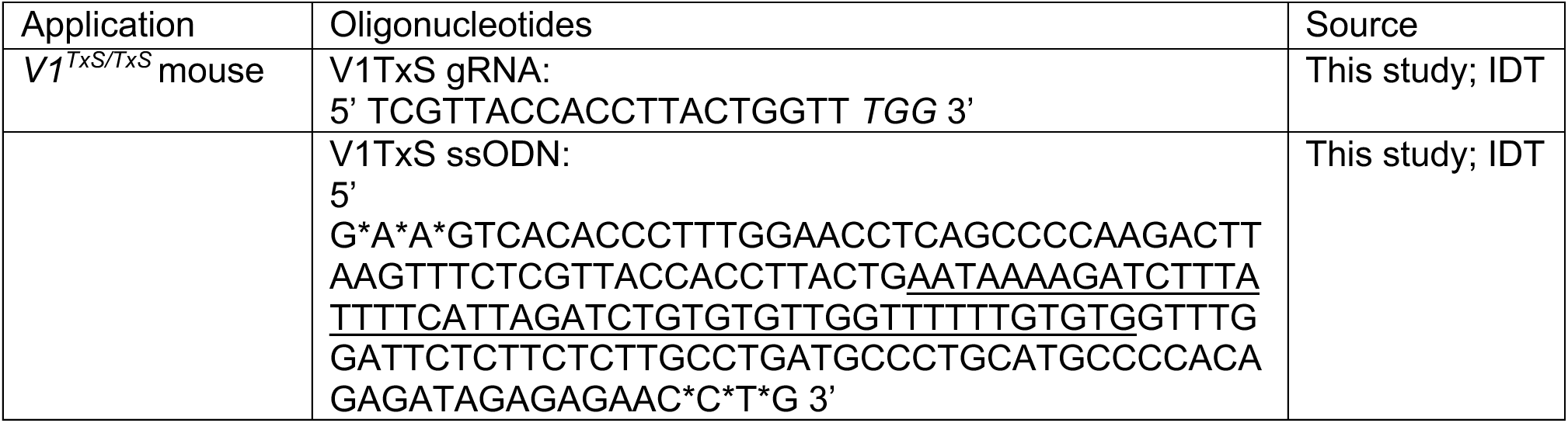
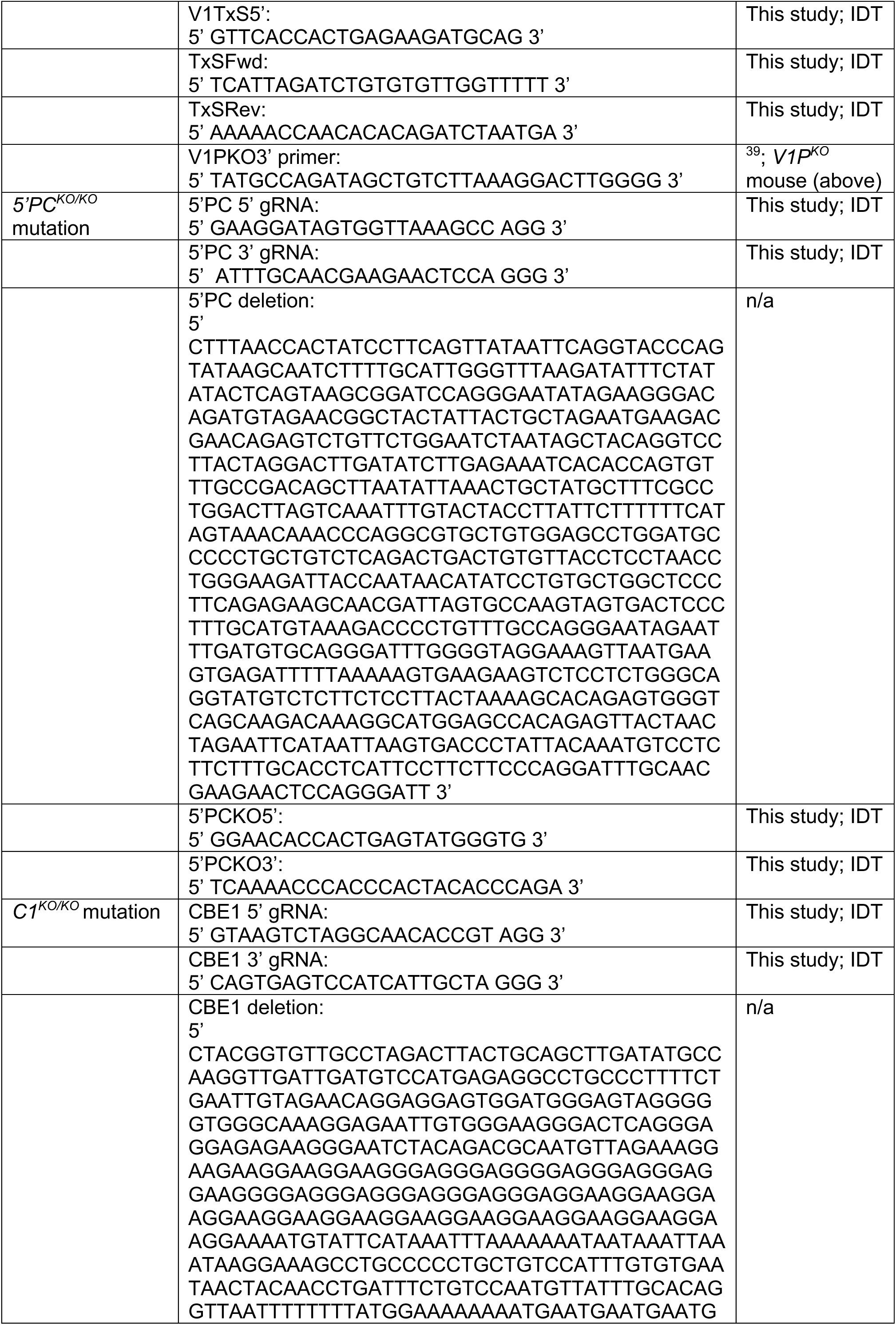
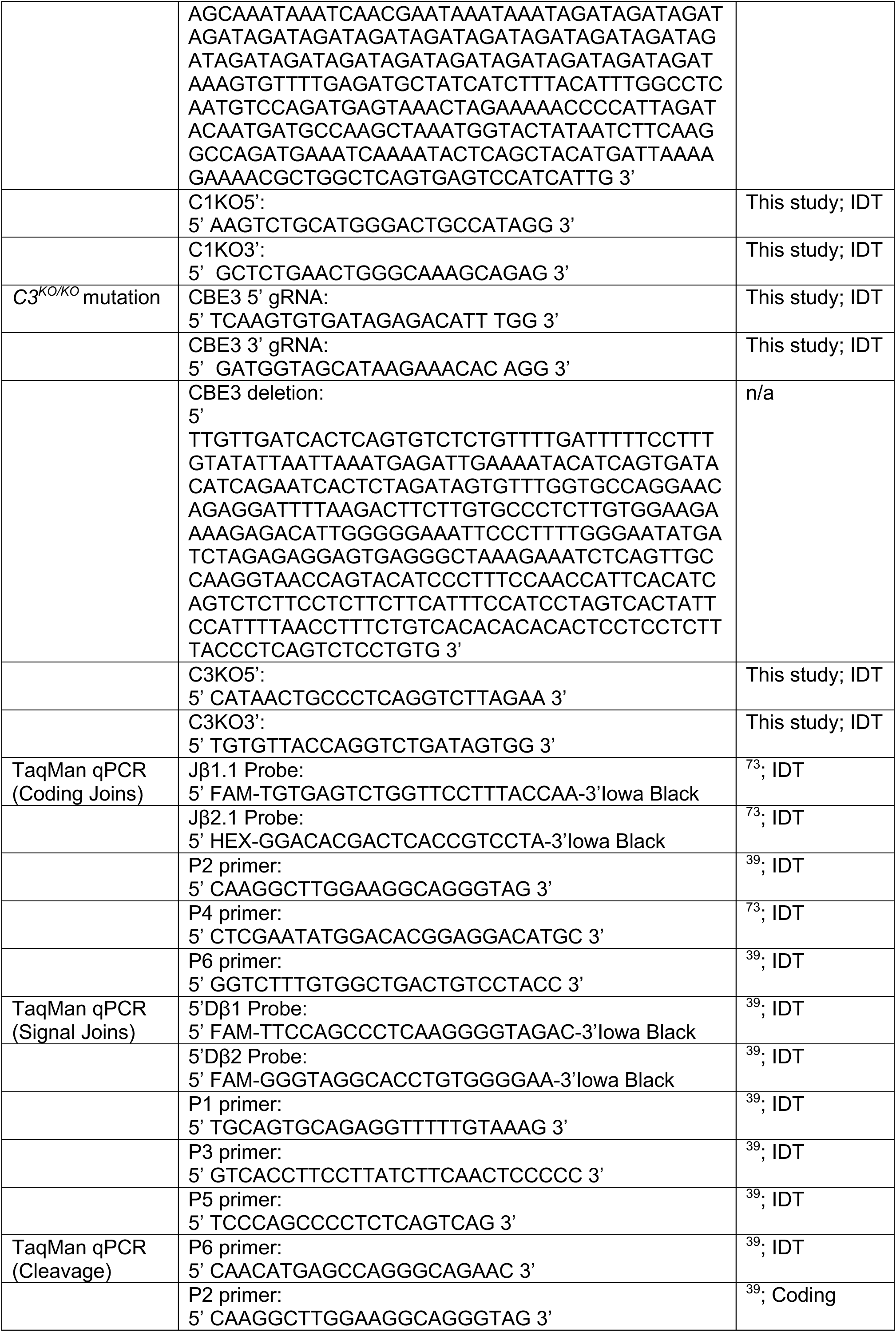
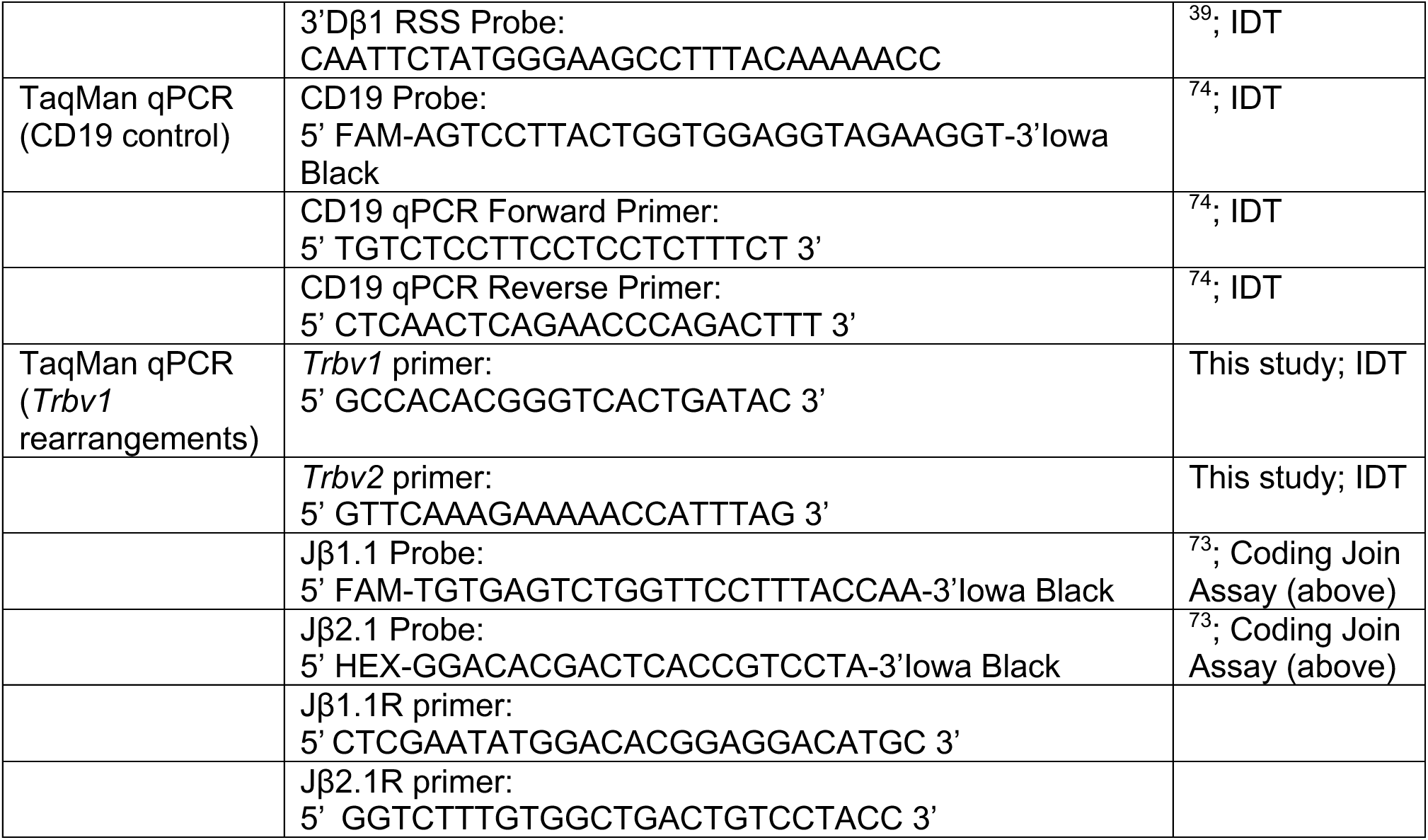
List of oligonucleotides.

### Real-time Quantitative PCR (qPCR)

TaqMan qPCR was performed with PrimeTime Gene Expression Master Mix (IDT,1055772) on sorted DN3 thymocytes (CD4^-^, CD8^-^, B220^-^, CD19^-^, CD11b^-^, CD11c^-^, NK1.1^-^, TER119^-^, TCRβ^-^, TCRγ/δ^-^, CD44^-^, CD25^+^) or total thymocytes from two RAG-sufficient mice per experiment (see Table 2 for complete list of antibodies). Primer/probe placement to assay for rearrangements was designed to amplify coding joins resulting from rearrangement to either DJβ1.1 or DJβ2.1 and either the *Trbv1* or *Trbv2* gene segment (See Table 1 for complete list of oligos). For statistical analysis we performed unpaired t tests using the Holm-Sidak method (α<0.05).

**Table 2.** List of reagents.

| Application | Antibody | Source | Catalog number |
| --- | --- | --- | --- |
| ChIP | Recombinant Anti-CTCF (Clone<br>EPR7314[B]) - ChIP Grade | Abcam | Cat# ab128873 |
| ChIP/CUT&<br>RUN | Anti-Histone H3 (acetyl K27) (polyclonal)-<br>ChIP Grade | Abcam | Cat# ab4729 |
| ChIP | Anti-Histone H3 (di methyl K9) (Clone<br>mAbcam 1220) - ChIP Grade | Abcam | Cat# ab1220 |
| ChIP | Anti-Histone H3 (tri methyl K4) antibody -<br>ChIP Grade | Abcam | Cat# ab8580 |
| CUT&RUN | CTCF CUTANA™ CUT&RUN Antibody | EpiCypher | SKU: 13-2014 |
| CUT&RUN | Anti-Rad21 antibody | Abcam | ab154769 |
| CUT&RUN | Rabbit anti-NIPBL Antibody, Affinity Purified | Bethyl<br>Laboratories | Cat# A301-779A |
| CUT&RUN | RNA Pol II Recombinant Mouse Monoclonal<br>Antibody | ActivMotif | Cat# 91152 |
| CUT&RUN | Anti-RNA polymerase II CTD repeat<br>YSPTSPS (phospho S2) antibody | Abcam | Cat# ab5095 |
| CUT&Tag | Histone H3K9me2 antibody (pAb) | ActivMotif | Cat# 39041 |
| DN and<br>DN3 sorts | Anti-B220 PE (Clone RA3-6B2) | BD Biosciences | Cat# 553090 |
| Flow | Anti-CD4 APC-eFluor780 (Clone RM4-5) | eBioscience | Cat# 47-0042-82 |
| DN3 sort | Anti-CD4 V450 |  |  |
| DN3 sort | Anti-CD8 APC |  |  |
| Flow | Anti-CD8 Pacific Blue (Clone 53-6.7) | BD Biosciences | Cat# 558106 |
| DN and DN3 sorts | Anti-CD11b PE (Clone M1/70) | BD Biosciences | Cat# 553311 |
| DN and DN3 sorts | Anti-CD11c PE (Clone HL3) | BD Biosciences | Cat# 561044 |
| Flow | Anti-CD16/CD32 | BD Biosciences | Cat# 553141 |
| DN and DN3 sorts | Anti-CD19 PE (Clone 1D3) | BD Biosciences | Cat# 553786 |
| DN3 sort | Anti-CD25 PE-Cy7 (Clone PC61) | BD Biosciences | Cat# 552880 |
| DN3 sort | Anti-CD44 FITC () | BD Biosciences |  |
| DN and DN3 sorts | Anti-NK1.1 PE (Clone PK136) | BD Biosciences | Cat# 553165 |
| Flow | Anti-TCRβ APC (Clone H57-597) | BD Biosciences | Cat# 561080 |
| DN and DN3 sorts | Anti-TCRβ PE (Clone H57-597) | BD Biosciences | Cat# 561081 |
| DN and DN3 sorts | Anti-TCR$\gamma/\delta$ Chain PE (Clone GL3) | BD Biosciences | Cat# 553178 |
| DN and DN3 sorts | Anti-TER-119 PE (Clone TER-119) | BD Biosciences | Cat# 553673 |
| Flow | Anti-Vβ2 (TRBV1) FITC (Clone B20.6) | BD Biosciences | Cat# 553280 |
| GRO Seq | BrdU Antibody (IIB5) AC | Santa Cruz Biotechnologies | Cat# sc-32323 AC |

### Flow Cytometry

Flow cytometry was performed on thymocytes from individual mice. Single-cell suspensions were prepared and treated with red blood cell lysis buffer (140 mM NH_4_Cl; 17 mM Tris, pH 7.4). Fc receptors were blocked with anti-CD16/CD32 and antibodies were stained in PBS with 2% FBS and 2 mM EDTA. To determine any effect on gross αβ T cell development and expression of TCRβ chains utilizing the *Trbv1* gene segment, we stained with the following panel: CD4, CD8, TCRβ, Vβ2 (TRBV1), and live/dead aqua (Life Technologies, L34957). Data were collected on an LSR Fortessa and analyzed with FlowJo software. Single cells were gated on based on forward and side scatter. For statistical analyses, we performed unpaired t tests using the Holm-Sidak method (α<0.05).

### RNA-Seq

RNA-Seq was performed on sorted (B220^-^, CD19^-^, CD11b^-^, CD11c^-^, NK1.1^-^, TER119^-^, TCRβ^-^, TCRγ/δ^-^) DN thymocytes from two *Rag1^-/-^* mice per experiment (see Table 2 for the list of antibodies). RNA extraction, library preparations, sequencing reactions, and bioinformatics analysis were conducted at GENEWIZ, LLC. (South Plainfield, NJ, USA). Total RNA was extracted from frozen cell pellet samples using Qiagen Rneasy Plus Universal mini kit followed by Manufacturer’s instructions (Qiagen, Hilden, Germany). RNA samples were quantified using Qubit 2.0 Fluorometer (Life Technologies, Carlsbad, CA, USA) and RNA integrity was checked with 4200 TapeStation (Agilent Technologies, Palo Alto, CA, USA). RNA sequencing library preparation was prepared using Rib0-Zero rRNA Removal Kit and TruSeq Stranded Total RNA library Prep kit following manufacturer’s protocol (Illumina, RS-122-2101). Briefly, rRNA was depleted with Ribp-Zero rRNA Removal Kit. rRNA depleted RNAs were fragmented for 8 minutes at 94 °C. First strand and second strand cDNA were subsequently synthesized. The second strand of cDNA was marked by incorporating dUTP during the synthesis. cDNA fragments were adenylated at 3’ends, and indexed adapter was ligated to cDNA fragments. Limited cycle PCR was used for library enrichment. The incorporated dUTP in second strand cDNA quenched the amplification of second strand, which helped to preserve the strand specificity. Sequencing libraries were validated using DNA Analysis Screen Tape on the Agilent 2200 TapeStation (Agilent Technologies, Palo Alto, CA, USA), and quantified by using Qubit 2.0 Fluorometer (Invitrogen, Carlsbad, CA) as well as by quantitative PCR (Applied Biosystems, Carlsbad, CA, USA). The sequencing libraries were multiplexed and clustered on three lanes of a flowcell. After clustering, the flowcell was loaded on the Illumina HiSeq instrument according to manufacturer’s instructions. The samples were sequenced using a 2 x150 Pair-End (PE) High Output configuration. Image analysis and base calling were conducted by the HiSeq Control Software (HCS) on the HiSeq instrument. Raw sequence data (.bcl files) generated from Illumina HiSeq was converted into fastq files and de-multiplexed using Illumina bcl2fastq program version 2.17. One mismatch was allowed for index sequence identification.

### Gro-Seq

GRO-Seq was performed on sorted (B220^-^, CD19^-^, CD11b^-^, CD11c^-^, NK1.1^-^, TER119^-^, TCRβ^-^, TCRγ/δ^-^) DN thymocytes from 7-9 *Rag1^-/-^* mice per experiment (see Table 2 for the list of antibodies) as previously described ^69^ with the following alterations. For the NRO buffer, 750μM of ATP, GTP and Br-UTP; 4.5μM of CTP; and 1.65% of N-Laurylsarcosine was used consistent with another publication where DN thymocytes were used ^70^. Following the isolation of nascent RNA, the Illumina Stranded Total RNA Prep, Ligation with Ribo-Zero Plus Kit (20040525) with the IDT for Illumina RNA UD Indexes Set A (20040529) were used for library prep minus the ribosomal RNA depletion steps. Paired-end sequencing was performed on an Illumina NextSeq 1000/2000 (300 cycles) v3.

### CUT & RUN

CUT & RUN was performed on sorted (B220^-^, CD19^-^, CD11b^-^, CD11c^-^, NK1.1^-^, TER119^-^, TCRβ^-^, TCRγ/δ^-^) DN thymocytes from 2-3 *Rag1^-/-^*mice per experiment (see Table 2 for the list of antibodies) using the EpiCypher CUTANA ChIC/CUT&RUN Kit (SKU: 14-1048) and CUTANA CUT&RUN Library Prep Kit (SKU: 14-1001) per the manufacturer’s instructions with the recommended nuclei extraction prior to adding the ConA beads. Notably, per the recommendation of starting with extra cells when performing nuclei extraction, as well as for quality control, 750K cells were used per reaction. Antibodies are listed in table 2. Paired-end sequencing was performed on an Illumina NextSeq 1000/2000 (100 cycles) v3.

### Micro Capture C

Micro Capture C was performed on sorted (B220^-^, CD19^-^, CD11b^-^, CD11c^-^, NK1.1^-^, TER119^-^, TCRβ^-^, TCRγ/δ^-^) DN thymocytes from 5-7 *Rag1^-/-^*mice per experiment (see Table 2 for the list of antibodies) using the Arima Custom Capture-HiC+ Kit (A510008). Two independent experimental replicates were performed per genotype for *WT, V1^TxS^*, and *513^TKO/TKO^*. Our custom panel was designed to span the *Tcrb* locus; chr6:39,771,104-42,660,754(mm10). Paired-end sequencing was performed on an Illumina NextSeq 1000/2000 (300 cycles) v3.

### Adaptive Immunosequencing

Adaptive Immunosequencing was performed on sorted DN3 thymocytes (CD4^-^, CD8^-^, B220^-^, CD19^-^, CD11b^-^, CD11c^-^, NK1.1^-^, TER119^-^, TCRβ^-^, TCRγ/δ^-^, CD44^-^, CD25^+^) pooled from two mice per experiment. Genomic DNA was isolated using the DNeasy Blood and Tissue Kit (Qiagen, 69506) and submitted to Adaptive Biotechnologies for their Mouse TCRβ assay at the survey resolution. For statistical analyses, we performed multiple unpaired t-tests.

### HiC

Conventional HiC was performed as previously described ^8,39,71^. Briefly, 5 × 10^6^ formaldehyde-cross-linked DN cells were lysed with 250 μl of ice-cold HiC lysis buffer (10 mM Tris-HCl pH 8.0, 10 mM NaCl, 0.2% IGEPAL CA630) containing protease inhibitors (Roche) for 15 minutes on ice. Chromatin was digested at 37 °C for 6 hours with DpnII (100 U), ends were filled and marked with biotin using Klenow and ligated together with T4 DNA ligase. Following the reversal of crosslinking, DNA was fragmented on a Covaris E220 Evolution Sonicator and size-selected for 300–500 bp with AMPure XP Beads (Beckman Coulter). DNA ends were repaired with the NEBNext Ultra II DNA Library Prep Kit according to the manufacturer’s instructions using 1 μg of the HiC DNA. The adapter-ligated DNA was size-selected for 300–400 bp with AMPure XP beads and biotinylated DNA fragments were pulled down using MyOne Streptavidin T1 beads (Life Technologies). The final HiC library was generated with 5 PCR cycles using the NEBNext Ultra II DNA Library Prep Kit and NEBNext Dual Index primers (NEB) for Illumina sequencing. For comparison of *WT* and *Eβ^KO/KO^*, we combined two replicates of conventional HiC with two replicates of Arima HiC using the Arima-HiC Kit (A510008) according to manufacture protocol with the KAPA Hyper Prep Kit. We performed these HiC on cells pooled from at least five mice for each genotype. Paired-end sequencing for both HiC methods was performed on an Illumina NovaSeq 6000 (300 cycles).

### Computational Analyses

For HiC, paired end raw reads were mapped with bwa version 0.7.17-r1188 (https://pubmed.ncbi.nlm.nih.gov/19451168/), treating forward and reverse reads separately, as described and implemented with the HiCExplorer command tool version 3.6 (https://pubmed.ncbi.nlm.nih.gov/29335486/). For Figure 1, we combined experimental data from conventional and Arima HiC because they had sufficient pairwise spearman correlations. For Figures 3 and 4, we down-sampled data from the genotype with the most reads to be equivalent to the number of reads from the genotype with the least reads. Visualization was done via UCSC genome browser (https://pubmed.ncbi.nlm.nih.gov/12045153/), using the interact format, and juicebox (https://pubmed.ncbi.nlm.nih.gov/29428417/). MUSTACHE version 1.2.0 (https://pubmed.ncbi.nlm.nih.gov/32998764/) was used to identify significant looping at resolutions 10K, 5K, 3K, and 1K. (part of the HiCUP package) to generate files that could be further processed to .hic files for visualization and loop calling with MUSTACHE. The figures show the resulting heatmaps with vanilla coverage (VC) normalization.

Raw RNA-Seq and GRO-Seq reads were trimmed using bbduk as part of bbmap (version 38.92) (https://sourceforge.net/projects/bbmap/). Reads were then aligned to mm10 using STAR (version 2.7.9a) (https://pubmed.ncbi.nlm.nih.gov/23104886/). bamCoverage from the deeptools package (version 3.5.1) (https://pubmed.ncbi.nlm.nih.gov/27079975/) yielded the bigwig coverage files and was run using flags to separate the coverage tracks by stranded-ness and to normalize to counts per million (cpm). Normalized bigwig coverage files were uploaded to the UCSC genome browser (https://pubmed.ncbi.nlm.nih.gov/12045153/). Gene expression levels were quantified using TPMcalculator (version 0.0.3) (https://pubmed.ncbi.nlm.nih.gov/30379987/) with the GENCODE annotation for the mm10 genome (version M23_GRCm38.p6). For normalization and identification of significantly differentially expressed genes, the raw read counts were analyzed with the limma-voom package from Bioconductor (version 3.54.2)(Law et al., 2014).

For ChiP-Seq, C&R and paired-end reads were aligned to the mm10 genome using bowtie2 (version 2.4.5) (https://pubmed.ncbi.nlm.nih.gov/22388286/). Peak calling was done using the macs2 algorithm (version 2.2.7.1) (https://pubmed.ncbi.nlm.nih.gov/18798982/). Bigwig files for the visualization in the UCSC genome browser were yielded by the bamCoverage tool inside the deeptools package (version 3.5.1) (https://pubmed.ncbi.nlm.nih.gov/27079975/) applying the RPGC normalization. Finally, the bioconducter package chipqc (version 1.30.0) (https://pubmed.ncbi.nlm.nih.gov/24782889/) helped determine our samples had a RiP (Reads in Peaks) score of at least 4.

