## Supplemental Figure 1 for "Promoter- and Enhancer-Dependent Cohesin Loading Initiates Chromosome Looping to Fold *Tcrb* Loci for Long-Range Recombination"

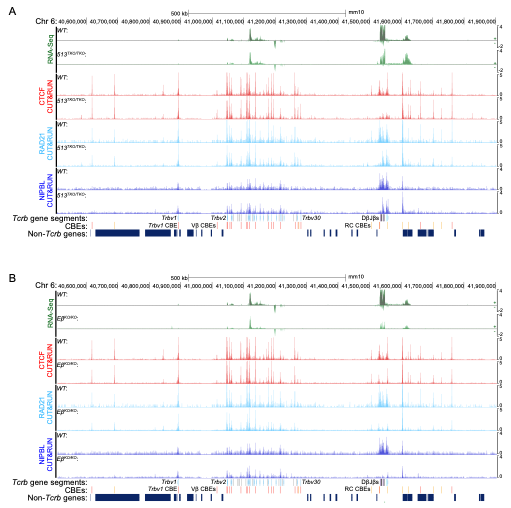

**Figure S1. . (A)** Genome browser tracks of *Tcrb* depicting two replicates each of RNA-Seq (sense and anti-sense transcripts) or a representative of two replicates of CUT&RUN for H3K27ac, CTCF, NIPBL, or RAD21 for *WT* or *513^TKO/TKO^* mice. **(B)** Genome browser tracks of *Tcrb* depicting two replicates each of RNA-Seq (sense and anti-sense transcripts) or a representative of two replicates of CUT&RUN for H3K27ac, CTCF, NIPBL, or RAD21 for *WT* or *Eβ^KO/KO^* mice.
